# 10-minimizers: a promising class of constant-space minimizers

**DOI:** 10.64898/2026.03.16.712052

**Authors:** Arseny Shur, Ido Tziony, Yaron Orenstein

## Abstract

Minimizers are sampling schemes which are ubiquitous in almost any high-throughput sequencing analysis. Assuming a fixed alphabet of size *σ*, a minimizer is defined by two positive integers *k, w* and a linear order *ρ* on *k*-mers. A sequence is processed by a sliding window algorithm that chooses in each window of length *w* + *k*− 1 its minimal *k*-mer with respect to *ρ* . A key characteristic of a minimizer is its density, which is the expected frequency of chosen *k*-mers among all *k*-mers in a random infinite *σ*-ary sequence. Minimizers of smaller density are preferred as they produce smaller samples, which lead to reduced runtime and memory usage in downstream applications. Recent studies developed methods to generate minimizers with optimal and near-optimal densities, but they require to explicitly store *k*-mer ranks in *Ω*(2^*k*^) space. While constant-space minimizers exist, and some of them are proven to be asymptotically optimal, no constant-space minimizers was proven to guarantee lower density compared to a random minimizer in the non-asymptotic regime, and many minimizer schemes suffer from long *k*-mer key-retrieval times due to complex computation.

In this paper, we introduce 10-minimizers, which constitute a class of minimizers with promising properties. First, we prove that for every *k* > 1 and every *w*≥ *k*− 2, a random 10-minimizer has, on expectation, lower density than a random minimizer. This is the first provable guarantee for a class of minimizers in the non-asymptotic regime. Second, we present spacers, which are particular 10-minimizers combining three desirable properties: they are constant-space, low-density, and have small *k*-mer key-retrieval time. In terms of density, spacers are competitive to the best known constant-space minimizers; in certain (*k, w*) regimes they achieve the lowest density among all known (not necessarily constant-space) minimizers. Notably, we are the first to benchmark constant-space minimizers in the time spent for *k*-mer key retrieval, which is the most fundamental operation in many minimizers-based methods. Our empirical results show that spacers can retrieve *k*-mer keys in competitive time (a few seconds per genome-size sequence, which is less than required by random minimizers), for all practical values of *k* and *w*. We expect 10-minimizers to improve minimizers-based methods, especially those using large window sizes. We also propose the *k*-mer key-retrieval benchmark as a standard objective for any new minimizer scheme.

## 1 Introduction

Sampling short substrings, termed *k*-mers, in long DNA sequences is a critical step in solving many bioinformatics tasks. Typically, a “window guarantee” is required: each window of *w* consecutive *k*-mers (i.e., a window of length *w* + *k*− 1) in the input sequence should be represented by at least one *k*-mer in the sample. Minimizers, introduced in [15,17], are simple sampling schemes with the window guarantee: a linear order on *k*-mers is fixed, and in each window, the minimal (lowest-ranked) *k*-mer w.r.t. this order is selected, with ties broken to the left. Minimizers are the most popular *k*-mer sampling schemes (a comprehensive list of bioinformatics applications using minimizers can be found in [12]), and even more advanced sampling schemes still use minimizers as an intermediate step [2,8,16].

Minimizers are most commonly characterized by their density, which is the expected fraction of sampled *k*-mers in an infinite sequence of i.i.d. symbols, as lower density leads to reduced runtime and memory usage of downstream applications. The average density (a.k.a. the expected density of a random minimizer) is close to 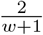 unless *w* ≫ *k* [5,17]. Many methods were designed to generate low-density minimizers. Methods like DOCKS [13], PASHA [3], and GreedyMini [6], generate orders explicitly, which highly limits the *k* values that can be used due to both the exponential, in *k* + *w*, construction time and *Θ*(*σ*^*k*^) space occupied by the order itself. (GreedyMini works with binary projections of DNA orders, which require *O*(2^*k*^) space, but this is still restrictive.)

Methods like miniception [18], double-decycling [14], and open-closed syncmers [13], define minimizer orders via *k*-mer comparison rules, without keeping the orders themselves. The obtained minimizers thus require only constant space to represent them. The property of being constant-space is desired for minimizers, as it enables the application of the minimizers to any value of *k*. Still, the existing constant-space minimizers suffer from two limitations. First, while these minimizers were shown to achieve low density empirically, no minimizer was provably guaranteed to have lower density than a random minimizer in the non-asymptotic regime. Second, *k*-mer comparison in constant-space minimizers can be viewed as computing a key for each *k*-mer and comparing the keys. For example, for a hash-based random minimizer, the key of the *k*-mer is its hash value, while for the double-decycling minimizer the primary key is the “class” of the *k*-mer, and the secondary key is the hash value. So far, no evaluation and benchmarking was performed on the *k*-mer key-retrieval time, which is a fundamental step in many bioinformatic application using minimizers. Some constant-space minimizers may suffer from complex computation, eventually slowing down an application that use them despite the low density they achieve.

In this paper, we present 10-minimizers, a novel class of minimizers with the first provable guarantee of achieving density better than random in the non-asymptotic regime. We prove that random 10-minimizers have the expected density 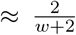 compared to 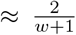 achieved by random minimizers. We present a particular family of constant-space 10-minimizers, called *spacers*, that demonstrate density results competitive to those of the known advanced constant-space minimizers. Moreover, for some (*k, w*) pairs, the spacers achieve the lowest density compared to all known (not necessarily constant-space) minimizers. Last, we formally define a performance-evaluation criterion for minimizers: *k*-mer key-retrieval time, and demonstrate that spacers are faster than hash-based random minimizers and most of competing constant-space minimizers. We expect 10-minimizers to improve the runtime and memory usage of many high-throughput sequencing methods by a simple replacement of their current *k*-mer key calculations.

## 2 Preliminaries

In what follows, Σ, σ, *k*, and *w* denote, respectively, the alphabet {0, *…, σ*− 1 }, its size, the length of the sampled substrings (*k*-mers), and the number of *k*-mers in a window. We write *s*[1..*n*] for a length-*n* string, denote the length of *s* by |*s*|, and use standard definitions of substring, prefix, and suffix of a string. The empty string is denoted by *ε*. We write *n*-string (*n*-prefix, *n*-suffix, *n*-window) to indicate length. We use the notation [*i*..*j*] for the range of integers from *i* to *j*, and *s*[*i*..*j*] for the substring of *s* covering the range [*i*..*j*] of positions.

Let *ρ* be a permutation (= a linear order) of Σ^*k*^. We view *ρ* as a bijection *ρ* : [1..*σ*^*k*^] → Σ^*k*^, binding all *k*-mers to their *ρ-ranks*. The *minimizer* (*ρ, w*) is a map *f* : Σ^*w* + *k*−1^ →[1..*w*] assigning to each (*w* + *k*−1)-window the starting position of its minimum- *ρ* -rank *k*-mer, with ties broken to the left. This map acts on strings (which we interchangeably call sequences) over Σ, selecting one position in each window so that in a window *v* = *S*[*i*..*i* + *w* + *k*− 2] the position *i* + *f* (*v*) −1 in *S* is selected.

Minimizers are evaluated by the density of selected positions. Let f(S) denote the set of positions selected in a string S by a minimizer f. The density of f is the limit 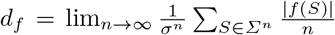 (also termed the expected density, to distinguish from the particular density over a specific sequence) Studying minimizer densities over a range of *w* values, it is convenient to “normalize” density to the *density factor D*_*f*_ = (*w* + 1)*d*_*f*_ .

An *arrangement π* of Σ^*k*^ with the *domain* ∅ = *U*⊆ Σ^*k*^ is an arbitrary permutation of *U* . In particular, arrangements with the domain Σ^*k*^ are exactly (linear) orders. We view *π* as a bijection *π* : [1.. |*U*|]→*U*, and apply the notion of *π*-rank to arbitrary arrangement *π*. We also consider *π* as an ordered list of length |*π*| =| *U*|, write *U*_*π*_ for the domain of *π* and *π*_1_.*π*_2_ for the arrangement obtained by concatenating arrangements *π*_1_ and *π*_2_ with disjoint domains, i.e., *U*_*π1*_ ∩*U*_*π2*_= ∅.

A subset *H* ⊆ Σ^*k*^ is a *universal hitting set* (UHS) *for w* if every (*w* + *k* −1)-window contains at least one *k*-mer from *H*. For example, {00, 01, 11} is a UHS for *σ*= *k* = 2 and any *w >* 1. We call an arrangement *π* a *UHS order for w* if *U*_*π*_ is a UHS for *w*. Then, any two orders *π*·*π*_1_ and *π*·*π*_2_ define the same minimizer for *w*, because the minimum-rank *k*-mer in every window is in *π*; we denote this minimizer by (*π, w*).

For a given minimizer *f* = (*ρ, w*), a (*w* + *k*)-window *v* (which contains two consecutive (*w* + *k*− 1)-windows and often termed *context*) is *charged* if its minimum-rank *k*-mer is either its prefix or its *unique suffix* (i.e., the *k*-suffix of *v* having no other occurrence in *v*); otherwise, *v* is *free*. Every string *S* contains exactly | *f* (*S*) | −1 charged (*w* + *k*)-windows [19, Lemma 6]. Since all possible *n*-strings have, in total, the same number of occurrences of each (*w* + *k*)-window, the density *d*_*f*_ of a minimizer equals the fraction of charged windows in Σ^*w* + *k*^ [11,18].

For an arrangement *π* of Σ^*k*^, a (*w* + *k*)-window *v* is *charged by π (due to π*[*i*]) if *π*[*i*] is the *k*-mer of minimal *π*-rank in *v*, and is a prefix (*prefix-charged*) or a unique suffix (*suffix-charged*) of *v*. The number of windows over Σ^*w* + *k*^ charged by *π* is denoted by ch_*w*_(*π*). A *live* window (w.r.t. a set *U* ⊂Σ^*w* + *k*^) is a window with no *k*-mer in *U* . If *U*⊂ Σ^*k*^ is disjoint from *U* _*π*_, then ch _*w*_(*U, π*) denotes the set of windows that are live w.r.t. *U* and charged by *π*. We say that an order or UHS order *ρ* is *initiated by U* if some prefix of *ρ* has the domain *U* .

Let (*ρ, w*) be a binary minimizer, *σ>* 2, and let *h* : Σ 0, 1 be a surjective *projection* map, extended to all strings over Σ letter-wise. A minimizer (*ρ* ^*′*^, *w*) over Σ is a *σ-extension of* (*ρ, w*) *with the projection h* if for any *k*-mers *u, v* ∈Σ^*k*^ the condition rank _*ρ*_ (*h*(*u*)) < rank _*ρ*_ (*h*(*v*)) implies rank _*ρ*_ *′* (*u*) < rank _*ρ*_ *′* (*v*) and rank _*ρ*_ (*h*(*u*)) *>* rank _*ρ*_ (*h*(*v*)) implies rank _*ρ*_ *′* (*u*) *>* rank _*ρ*_ *′* (*v*). In other words, *u* and *v* can be compared w.r.t. *ρ* ^*′*^ by comparing the binary *k*-mers *h*(*u*) and *h*(*v*) w.r.t. *ρ*, using *ρ* ^*′*^only to break ties.

In a similar way, *k*-extensions of arbitrary minimizers are defined. Let (*ρ, w*) and (*ρ* ^*′*^, *w*) be two minimizers such that *ρ* is an order on Σ^*k*^, *ρ* ^*′*^ is and order on 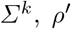, and *k*^*′*^ *> k*. If for any *k*^*′*^-mers 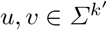 the condition rank _*ρ*_ (*u*[1..*k*]) < rank _*ρ*_ (*v*[1..*k*]) implies rank _*ρ*_ *′* (*u*) < rank _*ρ*_ *′* (*v*) and rank _*ρ*_ (*u*[1..*k*]) *>*rank _*ρ*_ (*v*[1..*k*]) implies rank _*ρ*_ *′* (*u*) *>* rank _*ρ*_ *′* (*v*), then (*ρ* ^*′*^, *w*) is a *k-extension* of (*ρ, w*).

We use Iverson notation: [expression] evaluates to 1/0 depending on expression being true/false.

## 3 Methods

### 3.1 10-minimizers

We start with the case Σ ={ 0, 1} . Let IO _*k*_ = {10*u*| *u∈* {0, 1} ^*k*−2^}; the elements of IO _*k*_ are *10-k-mers*. We consider the binary (UHS) orders initiated by IO _*k*_. First we solve the problem common to all such orders: to find an arrangement *τ* that hits all windows that are live w.r.t. IO _*k*_ and charges only few of them.

#### Lemma 1.

*Suppose that σ*= 2, *w* ≥ *k* − 2, *τ* = (1^*k*−1^0, 01^*k*−2^0, *…*, 0^*k*−2^10, 0^*k*−1^1, 1^*k*^, 0^*k*^), *and π is an arrangement of* IO _*k*_. *Then π · τ is a UHS for w and* ch _*w*_(IO _*k*_, *τ*) = *w* + *k* + 3 + [*w* = *k* − 2].

#### Proof.

Since a window is live w.r.t. IO _*k*_ if and only if it contains no *k*-mers beginning with 10, the set *X* of such live windows is given by *X* = {0^*i*^1^*w* + 2 −*i*^*u*| *i ∈* [0..*w* +2], *u* ∈{0, 1} ^*k*−2^ }. For 1≤ *i* ≤ *k* −1, any window with the prefix *τ* [*i*] = 0^*i*−1^1^*k*−*i*^0 does not belong to *X*: the longest prefix of the form 0^*i*^1^*j*^ in a window beginning with *τ* [*i*] has length *k* − 1, while the windows in *X* have such a prefix of length at least *w* + 2 *> k* − 1. Hence, *π · τ* prefix-charges 0 windows due to *τ* [1], *…, τ* [*k*−1]. As a unique suffix, 1^*k*−1^0 occurs in all live windows of the form 0^*i*^1^*w*−*i*^1^*k*−1^0, *i ∈* [0..*w*], so it suffix-charges *w* + 1 windows. Each of the *k*-mers 0^*i*^1^*k*−*i*−1^0, *i ∈* [1..*k*−2], is a unique suffix of exactly one window from *X*, namely, 0^*w* + *i*−1^1^*k*−*i*^0. Since a window from *X* cannot contain both *τ* [*i*] and *τ* [*j*], where 1 ≤ *i* < *j* ≤ *k* − 1, each *τ* [*i*], *i* ∈ [2..*k* − 1], suffix-charges one window.

Suppose that an *n*-window *v* is live w.r.t. *π · τ* [1..*k*−1] and contains 10. We can write *v* = *v* _1_10*v* _2_for some *v* _1_, *v* _2_ ∈ Σ^*∗*^ such that *v* _1_ does not contain 10. Then, |*v* _1_| ≤ *k* − 3 (otherwise, *v* contains some *τ* [*i*]) and |*v* _2_| ≤ *k* − 3 (otherwise, *v* contains a *k*-mer from *π*). Therefore, *n* ≤ 2*k* − 4 ≤ *w* + *k* − 2.

Hence, the live (*w* + *k*)-windows w.r.t. *π · τ* [1..*k* −1] form the set *X*^*′*^ = 0^*i*^1^*w* + *k*−*i*^ |*i* ∈ [0..*w* + *k*] (and for (*w* + *k* −1)-windows, the structure of the set is the same). Every window from *X*^*′*^ contain at least one of the *k*-mers *τ* [*k*] = 0^*k*−1^1, *τ* [*k* +1] = 1^*k*^, *τ* [*k* +2] = 0^*k*^, implying that *π·τ* is a UHS for *w*. It remains to note that *τ* [*k*] = 0^*k*−1^1 charges the windows 0^*k*−1^1^*w* +1^ and 0^*w* + *k*−1^1, *τ* [*k* +1] = 1^*k*^ charges 1^*w* + *k*^ and 0^*w*^1^*k*^ (the latter one only if it does not contain 0^*k*−1^1, i.e., for *w* = *k* − 2), and *τ* [*k* +2] = 0^*k*^ charges 0^*w* + *k*^. Summing up all windows charged due to *τ*, we get the amount stated in the lemma.

#### Remark 1.

The only *k*-mers from *τ* that can occur more than once in a live window w.r.t. IO _*k*_ are 1^*k*^ and 0^*k*^.

#### Definition 1.

*An order on* {0, 1 }^*k*^ *is a* 10-order *if it has the prefix π· τ, where π is an arbitrary arrangement of* IO _*k*_ *and τ is the arrangement defined in Lemma 1. An order *ρ* on* {0, …, *σ*−1}^*k*^ *is a* 10-order *if it is a σ-extension of a binary 10-order with a projection h* : {0, …, *σ* −1} → {0, 1} . *If, moreover*, |*h*^−1^(0) | =|*h*^−1^(1) |, *then *ρ* is a* balanced *10-order. A minimizer defined by a 10-order is called a* 10-minimizer.

### 3.2 Expected density of random 10-minimizers

#### Definition 2.

*A* random minimizer *is a random variable equidistributed over all minimizers with given parameters σ, k, and w. A* random 10-minimizer *is a random variable equidistributed over all 10-minimizers with given parameters σ, k, and w, and with a given projection h in the case σ >* 2.

The expected density of a random minimizer equals 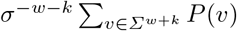, where *P* (*v*) is the probability of the window *v* being charged [5]. To estimate this expected density, a simplifying assumption was used [11], yielding an error that is small for wide ranges of parameters. Let us call a window *repetitive* if it contains some *k*-mer at least twice, and *repetition-free* if it is not repetitive. Informally, the assumption says “ignore all repetitive (*w* + *k*)-windows”. As such windows are rare unless *w*≫*k*, the error introduced by such an assumption can be proved small. Since any repetition-free (*w* + *k*)-window *v* contains (*w* +1) distinct *k*-mers, a random minimizer prefix-charges *v* with probability 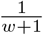 and suffix-charges *v* with the same probability. Hence, the simplifying assumption can be formalized directly as follows:

A. the random minimizer with the parameters (*σ, k, w*) prefix-charges any (*w* + *k*)-window with probability 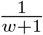 and suffix-charges it with the same probability 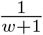 .

Assuming (A), one estimates the expectation of the density of a random (*σ, k, w*)-minimizer to 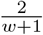 . Note that for a (*w* + *k*)-window with *t* distinct *k*-mers, the probability of being prefix-charged by a random minimizer is 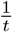, while the probability of being suffix-charged is either 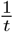 or 0, depending on the uniqueness of the *k*-suffix. Thus, for repetitive (*w* + *k*)-windows (A) underestimates the probability of being prefix-charged, while the probability of being suffix-charged can be affected in any direction.

In what follows, we formulate similar assumptions for random 10-minimizers and estimate their expected density based on these assumptions. We restrict ourselves to the case *w* ≥ *k* − 2. Therefore, *π · τ* is a UHS order for any arrangement *π* of IO _*k*_.

#### Case *σ* = 2

We introduce the simplifying assumption for random binary 10-minimizers, aiming to ignore only a fraction of all repetitive windows. Namely, a (*w* + *k*)-window *v* is *essentially repetitive* if there exists a *k*-mer *u* that occurs in *v* at least twice and is minimal in *v* in at least one 10-order but not in all 10-orders. Informally, the assumption says “ignore all essentially repetitive (*w* + *k*)-windows”.

The status (charged or free) of any window without 10-*k*-mers is the same for any 10-minimizer, and the number of charged windows of this type is computed in Lemma 1. Hence, essentially repetitive windows are those containing some 10-*k*-mer at least twice. If a (*w* + *k*)-window *v* contains *m* 10-*k*-mers, each occurring in *v* exactly once, then *v* is prefix-charged (respectively, suffix-charged) with probability 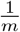 if its *k*-prefix (respectively, *k*-suffix) is a 10-*k*-mer. Accordingly, we assign the same probabilities to all (*w* + *k*)-windows with *m* occurrences of 10-*k*-mers, formalizing our assumption as follows:

(A^†^) the random 10-minimizer with the parameters (2, *k, w* ≥ *k*−2) prefix-charges (respectively, suffix-charges) any (*w* + *k*)-window *v* with *m* ≥ 1 occurrences of 10-*k*-mers with probability 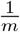 if *v* has a 10-*k*-mer as a prefix (respectively, suffix).

Using (A^†^), we get the following estimate, very close to 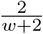, for the expected density ℛ (2, *k, w*) of a random binary 10-minimizer.

##### Theorem 1.

*Under the assumption* (A^†^), *the expected density* ℛ (2, *k, w*) *of a random 10-minimizer with the parameters* (2, *k, w* ≥ *k* − 2) *approximates to*

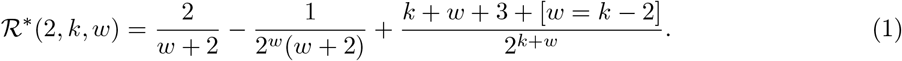

##### Proof.

Let *k* and *w* be fixed. For *m*≥ 1, we denote by IOP(*m*) (resp., IOS(*m*)) the set of (*w* + *k*)-windows with *m* occurrences of 10-*k*-mers, one of which is a prefix (resp., suffix) of the window. The function that deletes the first two letters of the *k*-suffix of a window and appends them to the beginning of the window is an obvious bijection from IOS(*m*) to IOP(*m*) for every *m* ≥ 1. Therefore, | IOP(*m*) | = | IOS(*m*) | . Assuming (A^†^) and taking Lemma 1 into account, we obtain

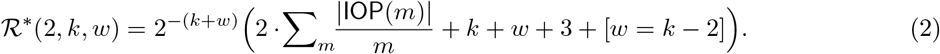

To compute |IOP(*m*)|, we note that a (*w* + *k*)-window *v* belongs to IOP(*m*) if and only if it *v* begins with 10 and contains *m* occurrences of 10 in the prefix *v*[1..*w* +2]. Equivalently,

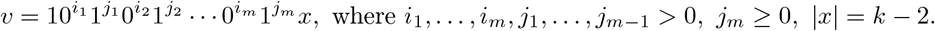

Respectively, the number of options for *v* is 2^*k*−2^ times the number of positive integer solutions to the inequality *i* _1_ + *j* _1_ + *· · ·* + *j* _*m*−1_ + *i* _*m*_ ≤ *w* + 1. This number of solutions is the number of possibilities to choose distinct values for the (2*m* − 1) partial sums

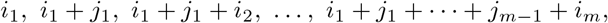

which is 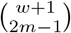 . Hence, 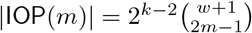 . To obtain (1), we prove a technical claim.

*Claim*. 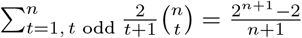 .

##### Proof.

We first observe that (i) 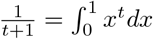 and thus we obtain (ii) 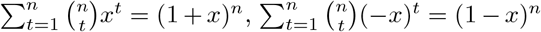 . As a result, we have

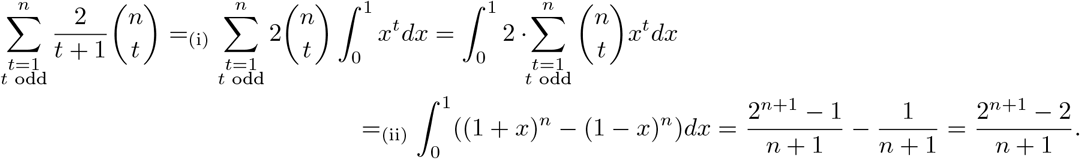

The claim is proved.

To finish the proof, we substitute the formula for |IOP(*m*)| into (2), using the Claim with *n* = *w* + 1, *t* = 2*m* − 1 to compute the sum (note that 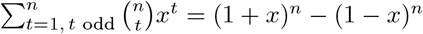) and finally obtain (1). □

#### Case σ ≥ 3

Let a 10-order *ρ* on Σ^*k*^ be the *σ*-extension of a binary 10-order *ρ* ^*^ by a projection *h*, and let *v* ∈ Σ^*k* + *w*^. If the binary window *h*(*v*) has a unique minimum *k*-mer in the order *ρ* ^*^, then *v* is charged by *ρ* if and only if *h*(*v*) is charged by *ρ* ^*^. Hence, the random 10-order with the parameters (*σ, k, w, h*) charges *v* with the same probability as the random 10-order with the parameters (2, *k, w*) charges *h*(*v*), except for two special cases of repetitive windows. If *h*(*v*) contains 10-*k*-mers, the special case arises if *h*(*v*) contains a repeated 10-*k*-mer; this case is exactly the one covered by the assumption (A^†^). If *h*(*v*) contains no 10-*k*-mers, its minimum-rank *k*-mer is from *τ* (Lemma 1), and hence is the same for all binary 10-orders. By Remark 1, 0^*k*^ and 1^*k*^ are the only *k*-mers in *τ* that can repeat in *h*(*v*), and one of these *k*-mers has the minimum rank in *h*(*v*) only if *h*(*v*) = 0^*w* + *k*^ or *h*(*v*) = 0^*i*^1^*w* + *k*−*i*^, where *i* ∈ [0..*k* −2]. This is the second special case where *v* and *h*(*v*) have different probabilities of being charged by the random 10-orders, but it affects a negligibly small set of windows (*k* distinct projections out of 2^*w* + *k*^ possible).

Thus, to be consistent with (A^†^), we state our assumption for the case *σ* ≥ 3 as follows:

(A^‡^) the random 10-minimizer with the parameters (*σ ≥* 3, *k, w*≥ *k* −2, *h*) charges any (*w* + *k*)-window *v* with the same probability as the random 10-minimizer with the parameters (2, *k, w*) charges the window *h*(*v*) under the assumption (A^†^).

Using (A^‡^), we extend Theorem 1 to *balanced* random 10-minimizers.

##### Theorem 2.

*Under the assumption* (A^‡^), *the expected density* ℛ(*σ, k, w, h*) *of a balanced random 10-minimizer with the parameters* (*σ ≥* 3, *k, w ≥ k*−2, *h*) *approximates to* ℛ ^*∗*^ (*σ, k, w, h*) = ℛ ^*∗*^ (2, *k, w*).

##### Proof.

For *v ∈Σ*^*w* + *k*^, let *P* (*v*) be the probability of *v* being charged by the random 10-minimizer with the parameters (*σ, k, w, h*), assigned according to (A^‡^). Similarly, *P* ^*′*^(*z*) denotes the probability that *z ∈* {0, 1}^*w* + *k*^ is charged by the random 10-minimizer with the parameters (2, *k, w*), assigned according to (A^†^). Then, 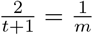 and 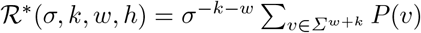 . Since 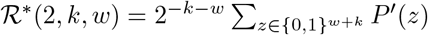, every window *z ∈* {0, 1}^*w* + *k*^ satisfies 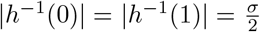 . By (A^‡^), all windows with the projection *z* have the same probability of being charged, equal to *P* ^*′*^(*z*). Then, 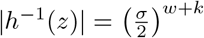 implying ℛ ^*∗*^ (*σ, k, w, h*) = ℛ ^*∗*^ (2, *k, w*). □

##### Remark 2.

The error of the approximations computed in Theorems 1, 2 can be roughly bounded by the fraction of (*w* + *k*)-windows with repeated *k*-mers among all (*w* + *k*)-windows. To evaluate the actual error, we computed the exact expected density in the case where *σ* = 2 and *k ∈* {7, 12}; see Section 4.1.

### 3.3 Lower density 10-minimizers

#### Existing 10-minimizers

In [1], the *ABB order* was proposed in the context of studying codes. This order compares any two *k*-mers over *Σ* using the alphabetic order for first letters and the rule 0 *>* 1 = 2 = *…* = *σ*−1 for subsequent letters. Neither [1] nor the later paper [4] (which applied this order as part of a minimizer) specified the tie-break used to make ABB a linear order. In [7], lexicographic tie-breaking was assumed and the resulting order, termed ABB+, was considered from the density point of view.

For every *k >* 1, the ABB+ order is an “almost” 10-order: it is obtained by *σ*-extension of a binary order *ρ* with the projection *h*(0) = 1, *h*(*a*) = 0 for *a* = 1, *…, σ −*1, and *ρ* is initiated by the set IO _*k*_. The sorting used to the rest of *k*-mers is inefficient compared to the arrangement *τ* employed in 10-orders (Lemma 1), but this makes a difference only for small *w* (for large *w*, the windows with no 10-*k*-mers in their projections are extremely rare). Since ABB+ in most cases has smaller density than the expected density of a random 10-minimizer, it serves as an important baseline for our minimizers.

#### Spacers

For a *k*-mer *u* ∈{0, 1} ^*k*^, its *tail* tail(*u*) is the longest proper suffix of *u* that is a prefix of a 10-*k*-mer. For example, tail(1001010) = 1010, tail(1001111) = 1, tail(1000000) = *ε*. To obtain binary 10-minimizers of lower density than expected by Theorem 1, we assign lower ranks to the 10-*k*-mers that ensure larger distance to the next 10-*k*-mer in the input sequence. In other words, we prioritize 10-*k*-mers with short tails.

##### Definition 3.

*The* tail score *of a 10-k-mer u is the ordered triple* (|tail(*u*) |, −bin(tail(*u*)), bin(*u*)) *of numeric keys, where* bin(*x*) *is the integer represented by x in binary. A* binary spacer (*ρ, w*) *is a binary 10-minimizer such that *ρ* ranks all 10-k-mers by their tail score, comparing the triples of keys lexicographically*.

##### Remark 3.

(1) The primary key |tail(*u*)| distinguishes binary spacers from binary ABB+ orders. For example, a binary spacer assigns the *k*-mer 101110 lower rank compared to 100100, while binary ABB+ (assuming 1 < 0) ranks these *k*-mers in reverse order.

(2)Using the inverse lexicographic order on tails as the secondary key showed the best empirical density results among the “simplest” tie-breakers.

(3)The choice of tertiary key has a very limited impact on the density (e.g., for *k* = 12 and 10 ≤*w* ≤36 choosing the inverse lexicographic order for the tertiary key instead of the lexicographic order changes the density by < 0.04% in all cases); nevertheless, the key is necessary to create a linear order.

##### Proposition 1.

*(1) Any binary spacer* (*ρ, w*) *is a constant-space minimizer. (2) Assuming that a k-mer fits into O*(1) *machine words, any two k-mers can be compared w*.*r*.*t. the order *ρ* in O*(1) *time*.

*Proof*. (1) Definition 3 gives an *O*(1)-size description of the binary spacer with any parameters (*k, w*). (2) It suffices to show that |tail(*u*)| can be found from the 10-*k*-mer *u* in *O*(1) operations. Viewing *u* as a binary integer, we compute the “indicator” ind(*u*) = *u* & ^*∼*^(*u* ≪ 1) (here &, ^*∼*^, and ≪ are the bitwise and, bitwise not, and left shift, respectively). The positions of 1’s in ind(*u*) are exactly the positions of 10’s in *u*. Then the second, from the left, occurrence of 1 in ind(*u*) is at the position of tail(*u*) in *u* by the definition of tail. Two more operations are needed to locate this position: a xor operation with the constant 2^*k*−1^ removes the leftmost 1, and the number of leading zeroes in the fixed-length representation of the obtained number is computed by the instruction lzcnt, which is standard for modern CPUs.□

Binary spacers have significantly lower density than binary ABB+. For example, for *w* = 24 and *k* ∈ [8..26], binary spacers have the density factors ≈1.74, compared to ≈1.86 for ABB+ and ≈1.92 for the random 10-minimizer according to Theorem 1 (Supplementary Table A2).

#### DNA spacers

As DNA minimizers are our primary interest, we consider 10-minimizers that are *σ*-extensions of binary spacers.

##### Definition 4.

*A* spacer *over an alphabet of size σ >* 2 *is a minimizer that is a σ-extension, with some projection h, of a binary spacer such that any k-mers u, v* ∈ Σ^*k*^ *with h*(*u*) = *h*(*v*) *are compared lexicographically*.

If a spacer is balanced (i.e., is a balanced 10-minimizer), then it inherits the density of its binary source almost exactly (for the details on the density guarantees for balanced *σ*-extensions see [6, Sect. 3.3]). On the other hand, *σ*-ary ABB+ minimizers, being *unbalanced σ*-extensions of binary minimizers, show much lower densities than in the binary case (density factor ≈1.72–1.73 for the above range *w* = 24, *k* ∈ [8..26], and the DNA alphabet). To reach even lower density, we focus on unbalanced spacers. For the 4-ary alphabet, we define the *DNA spacer* by the projection *h*(0) = *h*(1) = *h*(2) = 0, *h*(3) = 1.

##### Remark 4.

Given a sequence *x* of *n* bits, the subsequence of bits in even positions can be extracted from *x* in *O*(log *n*) bitwise operations by a standard FFT-like algorithm. As a DNA *k*-mer *u* is a sequence of 2*k* bits, its projection *h*(*u*) can be found in *O*(log *k*) operations by extracting the bits in even positions from the sequence (*u &* (*u* ≪ 1)) *≫* 1.

Proposition 1, Remark 4, and Definition 4 together imply

##### Proposition 2.

*(1) Any DNA spacer* (*ρ, w*) *is a constant-space minimizer*.

*(2) Assuming that a k-mer fits into O*(1) *machine words, any two k-mers can be compared w*.*r*.*t. the order *ρ* in O*(log *k*) *time*.

As is shown in Section 4.2, DNA spacers reach significantly lower density than ABB+ minimizers for the DNA alphabet.

### 3.4 Retrieving *k*-mer keys for DNA spacers

Fast sequence sampling with a minimizer (*ρ, w*) requires efficient solutions to two problems. First, the input sequence *S* should be converted into a sequence of fast-sortable keys, preferably numeric, representing all *k*-mers in *S*. Second, the minimum key in every set of *w* consecutive keys should be computed; this key represents the minimum *k*-mer w.r.t. *ρ* in the corresponding (*w* + *k* −1)-window. The second problem is independent of *ρ*, so we do not consider it here. Meanwhile, the solution to the first problem is *ρ* -specific unless *ρ* is explicitly stored. To be practically useful, a constant-space minimizer should be endowed with a fast key-retrieval algorithm. Let us formally define the problem.

#### Definition 5

(*k***-mer key-retrieval problem for a DNA minimizer)**.

Parameters: *σ* = 4, *k* ≥ 2, *w* ≥ 2, *an order ρ on* Σ^*k*^.

Input: *an arbitrary sequence S* = *S*[1..*n*] *∈ Σ ∗*.

Output: *a sequence K* = *K*[1..*n*−*k* +1] *of k*-mer keys, *satisfying the following property:*

(∗) *for every* (*w* + *k* − 1)*-window v* = *S*[*i*..*i* + *w* + *k*−2] *in S and every j* ∈ [*i*..*i* + *w*− 1], *the k-mer S*[*j*..*j* + *k*−1] *has the minimum *ρ* -rank among the k-mers in v if and only if K*[*j*] *is the minimum key among K*[*i*], *…, K*[*i* + *w*−1].

#### Remark 5.

The condition (∗) ensures that *K* suffices for a correct sampling of *S* with the minimizer (*ρ, w*). At the same time, (∗) gives a lot of freedom in assigning keys to *k*-mers. In particular, no bijection between *k*-mers and keys is required. Thus, if we learn that *S*[*j*..*j* + *k*−1] cannot be a minimum *k*-mer in any window it occurs in, then we can immediately set *K*[*j*] to be the “infinite” key.

Let us define the *k*-mer keys for DNA spacers. We call a *k*-mer *u 10-projected* if *h*(*u*) ∈ IO*k* and *τ* -projected if *h*(*u*) *∈ τ* .

- If the *k*-mer *u* is 10-projected, its key is the triple (|tail(*h*(*u*))|, − bin(tail(*h*(*u*))), bin(*u*));
- if *u* is *τ* -projected (namely, *h*(*u*) = *τ* [*i*]), the key of *u* is (*k, i*, bin(*u*));

- in both above cases, two first elements of the key are packed into one integer;

- if *u* is neither 10-projected nor *τ* -projected, its key is (*∞*, bin(*u*));
- a special key (*∞,∞*) is assigned to any *k*-mer that is found irrelevant to determine the minimum in all windows in which it appears.

#### Remark 6.

(i) by Definitions 3, 4, if *u* is 10-projected, then its key should include the tie-breaker bin(*h*(*u*)). Using Remark 3(iii), we drop bin(*h*(*u*)) from the key, as bin(*u*) suffices to break all ties.

i. Since |tail(*h*(*u*))| < *k*, the 10-projected *k*-mers are correctly compared against the *τ* -projected *k*-mers.
ii. Due to Lemma 1, tie breaks for the keys beginning with ∞ are redundant if *w* ≥ *k* − 2.

Now we describe the algorithm retrieving the *k*-mer keys for a DNA spacer.

1. The *k*-mers in the input string *S* are processed left to right;
2. for each *k*-mer, its projection is computed in *O*(1) time from the projection of the previous *k*-mer;
3. if the current *k*-mer *u* is 10-projected, its key is computed immediately; otherwise, *u* is pushed into the buffer *B* and gets its key later;
4. if the current *k*-mer *u* is 10-projected, the key (∞,∞) is assigned to all *k*-mers in *B*, and *B* is emptied;
5. *B* accommodates at most *w* −1 *k*-mers, all consecutive; if the *w*’th *k*-mer is pushed into *B*, the keys for all *k*-mers in *B* are computed by definition, and *B* is emptied; for all subsequent *k*-mers, the keys are computed immediately, until a 10-projected *k*-mer is encountered.

Step 4 is justified by Remark 5, as every window containing the *k*-mer to which the key (∞,∞) is assigned, contains at least one 10-projected *k*-mer. The expensive Step 5 is performed rarely unless *w* is small: for random *S*, the probability that the distance between two nearest 10-projected *k*-mers is at least *w* is *≈* 0.11 for *w* = 10 and < 0.0004 for *w* = 30.

## 4 Results

### 4.1 Estimated expected density vs. exact expected density for random 10-minimizers

To evaluate the actual error introduced by assumption (A^†^) and the closely related assumption (A^‡^) to the expected density of a random 10-minimizer, we designed an optimized enumeration algorithm (Supplementary Section A.1) that computes the *exact* expected density of a random 10-minimizer with the parameters (2, *k, w*) via the sum, over all (*w* + *k*)-windows containing 10-*k*-mers, of probabilities of being charged by such a random 10-minimizer. Recall that the number of charged windows among those containing no 10-*k*-mers is given by Lemma 1. For *k ∈* {7, 12}, *w∈* [*k* 2..2*k*], we compared the estimate (1) to the exact expected density (Supplementary Table A1). The error of the estimate is very small: < 0.11% for *k* = 7 and < 0.0022% for *k* = 12.

Thus, the estimate 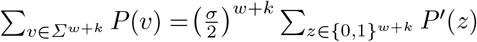 for the expected density of a random binary 10-minimizer is quite precise not only asymptotically but also for practical (*w, k*) ranges. This conclusion also extends to random balanced 10-minimizers over arbitrary even-size alphabets as the simplifying assumption (A^‡^) is based on (A^†^).

### 4.2 Density results for DNA spacers

We compared the density of DNA spacers to that of other constant-space minimizers, specifically: miniception [18], double-decycling [14], open-closed syncmers [13], and ABB+ [1,7]. We additionally included two baselines: the lowest density achieved by GreedyMini minimizers [6], which are not constant-space, and the KKMLT lower bound [10], which is valid for all *forward* schemes, including minimizers. We excluded mod-minimizer-based minimizers [9] from our comparisons as they are designed to work with *k > w* (even *k* ≥ *w* + 4 in the default setting), while DNA spacers are designed primarily for *w* ≥ *k*− 2. We measured the density of constant-space minimizers using a random DNA sequence of 2 *×*10^6^ nucleotides. We averaged the results of 5 runs over different sequences, ensuring to use different seeds for the minimizers that rely on randomness, i.e., miniception and open-closed-syncmer. For both miniception and open-closed-syncmer, which rank *s*-mers were *s* < *k*, we used *s* = 4. The density measurement for GreedyMini minimizers is detailed in Supplementary Section A.3.

We measured the density in the modes “fixed *k*, varying *w*” for *k ∈* {12, 24}, 3 ≤ *w* ≤ 100 (Fig. 1 AB), and “fixed *w*, varying *k*” for *w ∈* {12, 24} and 3 ≤ *k* ≤ 32 (Fig. 1 CD). For the DNA spacer, we used

**Fig. 1:**
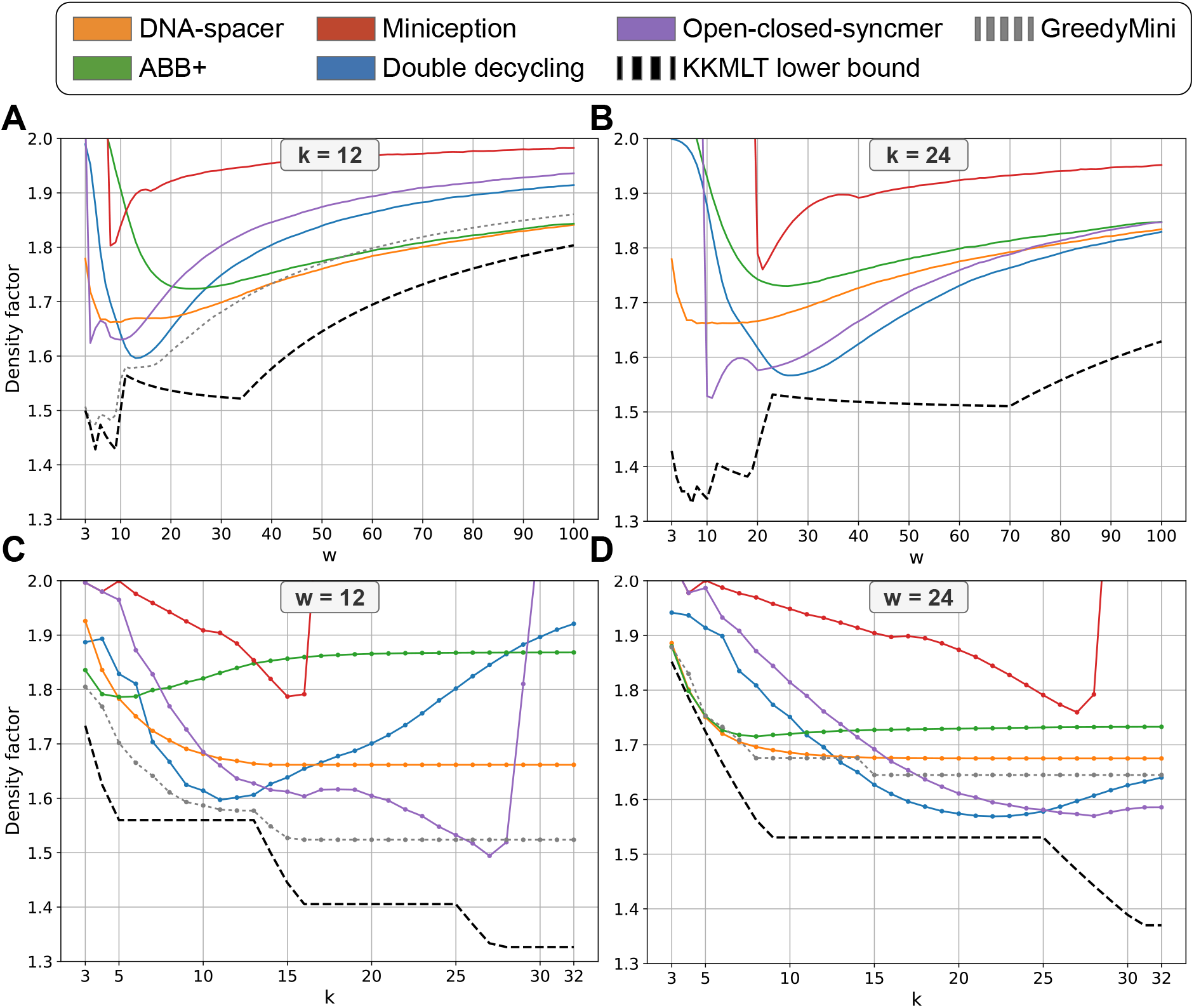
Performance of various DNA minimizers over certain ranges of (*k, w*) values. (A-B) Density comparisons over fixed *k ∈* {12, 24} and 3 ≤ *w* ≤ 100. (C-D) Density comparisons over fixed *w ∈* {12, 24} and 3 ≤ *k* ≤ 32.

*k*-extensions for the cases where *w* < *k* − 2. Namely, we compared the densities of the DNA spacer for (*k, w*) and the *k*-extension of the DNA spacer for (*w* +2, *w*) and lexicographic tie-breaking, and reported the lower density.

For *k* = 12, DNA spacer beats all other constant-space methods for *w* ≥ 23 and all methods, including GreedyMini, for *w* ≥ 40. For *k* = 24, double-decycling wins for *w* ≥ *k*, but DNA spacer catches it up as *w* grows to 100. In the fixed *w* mode, DNA spacer is still competitive, especially for *k* small enough. In particular, it beats all constant-space methods for *w* = 24, 5 ≤*k* ≤12, and is better than GreedyMini for *k* = 5, 6, 7.

### 4.3 Key-retrieval times for DNA spacers

The key-retrieval time of a minimizer heavily depends on the hardware, so the absolute figures may be insufficiently informative. To overcome this, we used two baseline hash-based minimizers: a trivial XOR-based hash and the C++ standard `std::hash`. We included in comparison DNA spacers (using the key-retrieval algorithm from Section 4.3), ABB+ minimizers (the algorithm is straightforward), and also miniception, open-closed syncmers and double-decycling. In the last three cases, we used our key-retrieval algorithms, as the open-closed-syncmer repository currently contains no sampling algorithm, and the miniception and double-decycling repositories present suboptimal sampling algorithms. We measured the processing time for a random DNA sequence of 1.5 *×* 10^8^ nucleotides, averaging the results over ten runs to ensure a robust estimate. All experiments were conducted on an AMD Ryzen 7800X3D system equipped with 32 GB of DDR5 RAM. The results of measurement for *w* = 24, 3 ≤ *k* ≤32 are shown in Fig. 2, A. DNA spacer beats all other minimizers except for ABB+ and the trivial hash; note the poor performance of open-closed syncmer and especially of double-decycling. In absolute figures, the DNA spacer’s key retrieval takes only several seconds for a genome-size input sequence. We also present Fig. 2, B to show that the key-retrieval times of DNA spacers do not increase as *w* grows. Higher times for *w* < 10 refer to steps 4 and 5 of the key-retrieval algorithm (Section 4.3): with a small buffer *B*, the expensive step 5 is performed more often.

**Fig. 2:**
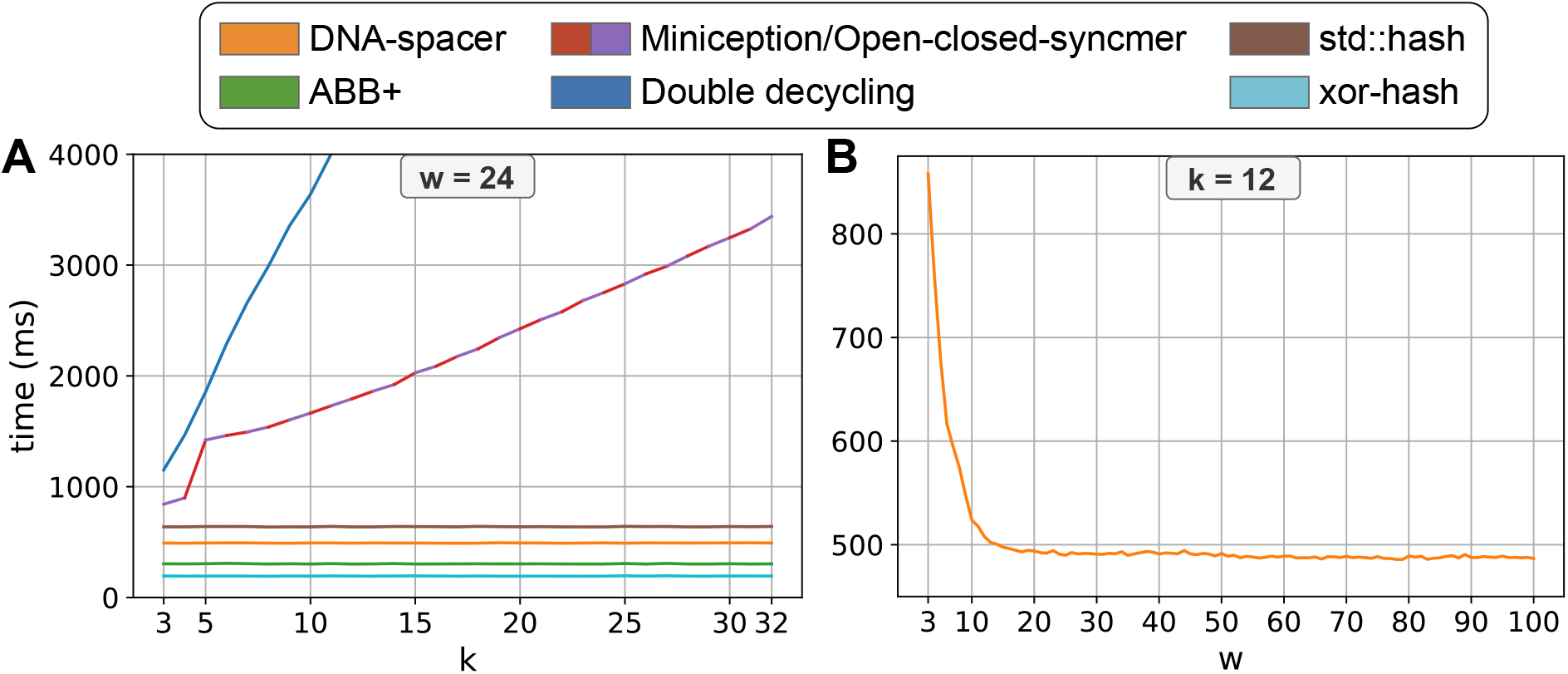
Time to retrieve keys for k-mers over a 1.5 10^8^ random DNA sequence, averaged over ten runs. Open-closed-syncmer times are plotted while miniception’s are not, as their implementations are very similar, which leads to less than 1% difference in most cases. (A) Key-retrieval times for *w* = 24 and 3 ≤ *k* ≤ 32. (B) Key-retrieval times for *k* = 12 and 3 ≤ *w* ≤ 100 (DNA spacers only).

## 5 Discussion

10-minimizers represent a significant advance in *k*-mer sampling schemes for high-throughput sequencing analysis, introducing the first class of constant-space minimizers with provable density guarantees superior to random minimizers in the non-asymptotic regime. By proving that random 10-minimizers achieve an expected density of approximately 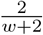 (vs. 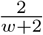 for random minimizers), we established a theoretical foundation for their superiority across practical *k, w* ranges, validated by exact density computations showing estimation errors < 0.0022% for *k* = 12.

Spacers, our flagship 10-minimizers, combine three desirable properties: (1) constant space (O(1) description, no explicit order storage); (2) low-density competitive with state-of-the-art schemes, i.e., achieving the lowest known density for certain (*k, w*) regimes (e.g., for 40 ≤*w* ≤100 and *k* = 12, or 5 ≤ *k* ≤ 7 and *w* = 24); and (3) fast *k*-mer rank retrieval, outperforming hash-based random minimizers.

Key insights from our analysis include: (1) maximization of inter-minimizer distances via short tails prioritization reduces the density; (2) 10-minimizers using unbalanced projections achieve lower density compared to those using balanced projections; (3) spacers are an attractive choice for sampling over large windows; and (4) key-retrieval time is an important metric for minimizer evaluation.

There are several open questions and future steps that are raised by our study. First, can we push the density of a constant-space minimizer further down, while keeping the key-retrieval time below the random minimizer baseline? Up to now, we were able to further improve the density of 10-minimizers only at the expense of more elaborate keys, which boost the key-retrieval time. Second, is it possible to obtain further theoretic density estimates, for example, on the expected density of a random unbalanced 10-minimizer or on the density of DNA spacer? Third, can we improve current high-throughput sequencing analysis methods and advanced sampling schemes by plugging in the best constant-space minimizer for their (*k, w*) values?

To conclude, 10-minimizers, particularly spacers, offer a theoretically-grounded, practically-efficient alternative to existing schemes, poised to enhance high-throughput sequencing pipelines through provably superior density and competitive retrieval times.

## Code and Supplementary Tables

https://github.com/OrensteinLab/10-minimizers

## Acknowledgment

Ido Tziony acknowledges the VATAT fellowship for excellent PhD students in Data Science, the Bar-Ilan University presidential fellowship, and the cloud-computing credit support of the Israel Data Science and AI Initiative.

## Funding

This study was supported by Israel Science Foundation grant no. 358/21 to Yaron Orenstein.

## Author contribution

A.S. performed the theoretical analysis and led the project, A.S. and I.T. developed, implemented, and benchmarked the algorithms, all authors analyzed the results and wrote the manuscript, and Y.O. supervised I.T. and acquired the funding.

## Notes

https://github.com/OrensteinLab/10-minimizers

### Competing Interest Statement

The authors have declared no competing interest.

## References

1. S. R. Blackburn, Non-overlapping codes, IEEE Transactions on Information Theory, 61 (2015), pp. 4890–4894.

2. R. Edgar, Syncmers are more sensitive than minimizers for selecting conserved k-mers in biological sequences, PeerJ, 9 (2021), p. e10805.

3. B. Ekim, B. Berger, and Y. Orenstein, A randomized parallel algorithm for efficiently finding near-optimal universal hitting sets, in International Conference on Research in Computational Molecular Biology, Springer, 2020, pp. 37–53.

4. M. C. Frith, L. Noé, and G. Kucherov, Minimally overlapping words for sequence similarity search, Bioinformatics, 36 (2020), pp. 5344–5350.

5. S. Golan and A. M. Shur, Expected density of random minimizers, in SOFSEM 2025: Theory and Practice of Computer Science - 50th International Conference on Current Trends in Theory and Practice of Computer Science, SOFSEM 2025, Proceedings, Part I, R. Královic and V. Kurková, eds., vol. 15538 of Lecture Notes in Computer Science, Springer, 2025, pp. 347–360.

6. S. Golan, I. Tziony, M. Kraus, Y. Orenstein, and A. Shur, GreedyMini: generating low-density DNA minimizers, Bioinformatics, 41 (2025), pp. i275–i284.

7. R. Groot Koerkamp, Low density minimizers, 2025.

8. R. Groot Koerkamp and G. E. Pibiri, The mod-minimizer: A simple and efficient sampling algorithm for long k-mers, in 24th International Workshop on Algorithms in Bioinformatics, WABI 2024, S. P. Pissis and W. Sung, eds., vol. 312 of LIPIcs, Schloss Dagstuhl - Leibniz-Zentrum für Informatik, 2024, pp. 11:1–11:23.

9. R. Groot Koerkamp and G. E. Pibiri, The mod-minimizer: A Simple and Efficient Sampling Algorithm for Long k-mers, in 24th International Workshop on Algorithms in Bioinformatics (WABI 2024), S. P. Pissis and W.-K. Sung, eds., vol. 312 of Leibniz International Proceedings in Informatics (LIPIcs), Dagstuhl, Germany, 2024, Schloss Dagstuhl – Leibniz-Zentrum für Informatik, pp. 11:1–11:23.

10. B. Kille, R. Groot Koerkamp, D. McAdams, A. Liu, and T. J. Treangen, A near-tight lower bound on the density of forward sampling schemes, Bioinformatics, 41 (2024), p. btae736.

11. G. Marçais, D. Pellow, D. Bork, Y. Orenstein, R. Shamir, and C. Kingsford, Improving the performance of minimizers and winnowing schemes, Bioinformatics, 33 (2017), pp. i110–i117.

12. M. Ndiaye, S. Prieto-Baños, L. M. Fitzgerald, A. Yazdizadeh Kharrazi, S. Oreshkov, C. Dessi-moz, F. J. Sedlazeck, N. Glover, and S. Majidian, When less is more: sketching with minimizers in genomics, Genome Biology, 25 (2024), p. 270.

13. Y. Orenstein, D. Pellow, G. Marçais, R. Shamir, and C. Kingsford, Designing small universal k-mer hitting sets for improved analysis of high-throughput sequencing, PLoS Computational Biology, 13 (2017), p. e1005777.

14. D. Pellow, L. Pu, B. Ekim, L. Kotlar, B. Berger, R. Shamir, and Y. Orenstein, Efficient mini-mizer orders for large values of k using minimum decycling sets, Genome Research, 33 (2023), pp. 1154–1161.

15. M. Roberts, W. Hayes, B. R. Hunt, S. M. Mount, and J. A. Yorke, Reducing storage requirements for biological sequence comparison, Bioinformatics, 20 (2004), pp. 3363–3369.

16. K. Sahlin, Effective sequence similarity detection with strobemers, Genome Research, 31 (2021), pp. 2080–2094.

17. S. Schleimer, D. S. Wilkerson, and A. Aiken, Winnowing: local algorithms for document fingerprinting, in Proceedings of the 2003 ACM SIGMOD International Conference on Management of Data, SIGMOD ‘03, New York, NY, USA, 2003, Association for Computing Machinery, p. 76–85.

18. H. Zheng, C. Kingsford, and G. Marçais, Improved design and analysis of practical minimizers, Bioinformatics, 36 (2020), pp. i119–i127.

19. H. Zheng, G. Marçais, and C. Kingsford, Creating and using minimizer sketches in computational genomics, Journal of Computational Biology, 30 (2023), pp. 1251–1276.

