## Appendix for "10-minimizers: a promising class of constant-space minimizers"

### A Appendix

#### A.1 Computing the exact density of random binary 10-minimizers

To compute the density of a random binary 10-minimizer with parameters $k, w$, it suffices to find, for each $(w+k)$-window containing at least one 10-$k$-mer, the probability of being charged (the case of windows with no 10-$k$-mers is covered by Lemma 1). For a $(w+k)$-window $v$ with $m$ distinct $k$-mers, this probability can be expressed as

$$\frac{[k\text{-prefix of } v \text{ is a 10-}k\text{-mer}] + [k\text{-suffix of } v \text{ is unique and is a 10-}k\text{-mer}]}{m}$$

The efficient enumeration of all binary $(w+k)$-windows is organized as follows.

1. The set of all $(w+k)$-windows is viewed as a binary *prefix tree*; its nodes are all binary strings of length $\leq w+k$, and its edges are all pairs $(u, ua)$, where $a \in \{0,1\}$. In particular, the leaves of this tree are exactly all the $(w+k)$-windows.
2. The prefix tree is processed by a recursive DFS; each node $za$ gets from its parent $z$ the set of 10-$k$-mers occurring in $z$ and the flag indicating whether the $k$-prefix of $z$ is a 10-$k$-mer. Hence, if $za$ is a leaf, the probability of $za$ being charged is computed in $O(1)$ operations.
3. To speed up the DFS traversal, the recursive routine is called for a node $z$ only if the $k$-suffix of $z$ is a 10-$k$-mer (and thus may change the set of occurred 10-$k$-mers inherited from the calling ancestor). If no descendants of a node $z$ have 10-$k$-mers as suffixes, then all leaves in the subtree of $z$ have the same probability of being charged; this probability is computed at $z$ and multiplied by the number of those leaves.

We computed the densities of random binary 10-minimizers with $k = 7$ and $k = 12$ over ranges of $w$ values and compared these results to the estimate $\mathcal{R}^*(2,k,w)$ by Theorem 1. The results show that the error of the estimate $\mathcal{R}^*(2,k,w)$ is less than 0.11% for $k = 7$ and less than 0.0022% for $k = 12$ (Table A1).

$k = 7$

| $w$ | 5 | 6 | 7 | 8 | 9 | 10 | 11 | 12 | 13 | 14 |
|---|---|---|---|---|---|---|---|---|---|---|
| $\mathcal{R}(2,7,w)$ | 1.711426 | 1.749573 | 1.780111 | 1.801437 | 1.820292 | 1.834556 | 1.848158 | 1.858888 | 1.868591 | 1.877106 |
| $\mathcal{R}^*(2,7,w)$ | 1.710938 | 1.750000 | 1.779134 | 1.801428 | 1.819310 | 1.834117 | 1.846664 | 1.857462 | 1.866860 | 1.875114 |

$k = 12$

| $w$ | 10 | 11 | 12 | 13 | 14 | 15 | 16 | ··· |
|---|---|---|---|---|---|---|---|---|
| $\mathcal{R}(2,12,w)$ | 1.832525 | 1.845776 | 1.856962 | 1.866604 | 1.874966 | 1.882349 | 1.888892 | ··· |
| $\mathcal{R}^*(2,12,w)$ | 1.832504 | 1.845739 | 1.856936 | 1.866564 | 1.874949 | 1.882328 | 1.888876 | ··· |

| ··· | 17 | 18 | 19 | 20 | 21 | 22 | 23 | 24 |
|---|---|---|---|---|---|---|---|---|
| ··· | 1.894742 | 1.900005 | 1.904763 | 1.909094 | 1.913039 | 1.916660 | 1.919991 | 1.923064 |
| ··· | 1.894731 | 1.899997 | 1.904760 | 1.909090 | 1.913043 | 1.916666 | 1.920000 | 1.923077 |

Table A1: Comparing exact ($\mathcal{R}(2,k,w)$) and estimated ($\mathcal{R}^*(2,k,w)$, Theorem 1) values of the expected density of random binary 10-minimizers.

#### A.2 Density of binary spacers vs. binary ABB+ minimizers

We compared the densities of binary spacers and binary ABB+ orders for $w = 24$ and $k = 8, \ldots, 26$, estimating them on a random sample of $10^8$ $(w+k)$-windows for each pair $(k, w)$ (Supplementary Table A2). We note that the estimate $\mathcal{R}^*(2,k,w)$ of the expected density factor of a random binary 10-minimizer, rounded to 6 decimal places, equals 1.923077 for all points in the analysed range.

$w = 24$

| $k$ | 8 | 9 | 10 | 11 | 12 | 13 | 14 | 15 | 16 | ··· |
|---|---|---|---|---|---|---|---|---|---|---|
| $D_{\mathsf{spacer}}(2,k,24)$ | 1.744185 | 1.741076 | 1.741098 | 1.742237 | 1.742177 | 1.742321 | 1.742793 | 1.743476 | 1.742896 | ··· |
| $D_{\mathsf{ABB+}}(2,k,24)$ | 1.854068 | 1.857927 | 1.858477 | 1.860609 | 1.858185 | 1.860894 | 1.860402 | 1.861672 | 1.861706 | ··· |

| ··· | 17 | 18 | 19 | 20 | 21 | 22 | 23 | 24 | 25 | 26 |
|---|---|---|---|---|---|---|---|---|---|---|
| ··· | 1.743550 | 1.743088 | 1.742920 | 1.742048 | 1.744163 | 1.741836 | 1.742611 | 1.742745 | 1.742599 | 1.743465 |
| ··· | 1.862441 | 1.861741 | 1.861548 | 1.860859 | 1.860694 | 1.859866 | 1.861647 | 1.860313 | 1.858494 | 1.859848 |

Table A2: Comparing the density factors of binary spacer and binary ABB+ minimizer. The densities are estimated from a sample of $10^8$ random $(w+k)$-windows for each pair $(k, w)$.

##### A.3 Calculating density of GreedyMini orders

To measure the density of GreedyMini orders, we conducted measurements using a random DNA sequence of $10^7$ nucleotides, or incorporated densities reported in the GreedyMini study [6]. The density values were measured as follows:

1. **For** $k = 12$ **and** $w \in [3, 100]$**:** We utilized the density values reported in the GreedyMini study. Additionally, we evaluated published GreedyMini orders for $(k, w) \in \{(12, 20), (12, 20)\}$ across smaller $w$ values. For each parameter set, we reported the minimum density observed between the original published results and our supplementary measurements.
2. **For** $k = 24$ **and** $w \in [3, 100]$**:** GreedyMini densities were omitted for this range, as this $k$ value exceeds the recommended $k$ range for GreedyMini.
3. **For** $w = 12$ **and** $k \in [3, 100]$**:** We utilized the densities reported in the GreedyMini study. For $k$ values not explicitly covered in the original publication, we estimated the density by applying the $k$-extension with lexicographic tie-breaking to the published GreedyMini order constructed for the largest available $k$ and $w = 12$.
4. **For** $w = 24$ **and** $k \in [3, 100]$**:** In the absence of published GreedyMini orders for these specific parameters, we calculated the density by applying the following orders (using $k$-extensions as needed) to measure the density for the setting of $(k, w) = (k', 24)$:
   - All $(k', 15)$ orders for $3 \le k' \le 14$.
   - The $(15, 17)$ orders for $15 \le k' \le 32$.
   - The $(15, 30)$ orders for $15 \le k' \le 32$.
   - The $(8, 19)$ orders for $8 \le k' \le 32$.

   For every $(k, w)$ configuration, we report the lowest density observed across all tested orders.
